# Time-varying habitat selection analysis: a model and applications for studying diel, seasonal and post-release changes

**DOI:** 10.1101/2023.01.04.522602

**Authors:** Romain Dejeante, Marion Valeix, Simon Chamaillé-Jammes

## Abstract

Resource selection functions are commonly employed to evaluate animals’ habitat selection, e.g. the disproportionate use of habitats relative to their availability. While environmental conditions or animal motivations may vary over time, sometimes in an unknown manner, studying changes in habitat selection usually requires *a priori* time discretization. This limits our ability to precisely answer the question ‘when is an animal’s habitat selection changing?’. Here, we present a straightforward and flexible alternative approach based on fitting dynamic logistic models to used/available data. First, using simulated dataset, we demonstrate that dynamic logistic models performed well to recover temporal variations in habitat selection. We then show real-world applications for studying diel, seasonal, and post-release changes in habitat selection of blue wildebeest (*Connochaetes taurinus*). Finally, we provide the relevant R scripts to facilitate the adoption of the method by ecologists. Dynamic logistic models allow to study temporal changes in habitat selection in a framework consistent with resource selection functions, but without the need to discretize time, which can be a difficult task when little is known about the process studied, or may obscure inter-individual variability in timing of change. These models should undoubtedly find their place in the movement ecology toolbox.

## 1. Introduction

Changes in environmental conditions consistently challenge animals in their lives, leading them to regularly adjust their behavior. One of the important way animals do so is by using the landscape they live in differently, i.e. by relocating themselves or selecting habitats differently. This may be most clearly exemplified by migrations, which occur in response to seasonal changes in weather and/or resource availability (Dingle 2014). Changes at smaller time-scales also occur, such as when animals shift habitats in response to forage depletion or day/night alternation in predation risk (Courbin et al. 2019). Naturally, discovering and understanding such changes in space use and habitat selection is a key goal of ecologists.

Over time, habitat selection analysis (HSA) conducted using the resource selection function (RSF) approach (Boyce et al. 2002) has become the standard framework to study changes in animals’ habitat selection. RSF analyses statistically compare the environmental characteristics of used locations collected over a period of time with the characteristics of locations available during that period. As such, RSF estimates the average strength of selection for the various habitats considered over the period of interest. How this period is defined is up to the researcher, but strongly affects the results and the associated interpretations (Mayor et al. 2009). As the within-period variability in selection is averaged, finer-scale temporal dynamics (e.g. day/night changes when the period covers weeks or months) in selection are overlooked, and the mean selection strength estimated might represent an average that is not meaningful. This would be the case if, for instance, the study period encompasses two different phases in an animal’s habitat selection behavior without the researcher being aware of it.

Discretizing time to define biologically relevant periods over which to conduct HSA may be difficult, and often involves somewhat arbitrary decisions with unknown consequences. This is true for even apparently obvious periods like seasons (Basille et al. 2013) or day-night periods (Richter et al. 2020). Starts and ends of seasons vary between years, and can only be roughly defined without ancillary data. Some seasons like spring or fall are also clearly periods of environmental changes during which patterns of habitat selection are unlikely to be constant. Animal needs and motivations, and thus habitat selection (Roever et al. 2014) may also change at unpredictable (for the researcher) times, such as when they disperse (Delgado et al. 2009). This again makes discretization of time difficult, or even irrelevant if one is interested in the dynamics of the change itself. This issue has been recognized before and various suggestions have been offered, from using a combination of movement metrics and habitat use information to define periods (but without estimating habitat selection) (Basille et al. 2013), to integrating time directly into the RSF predictor (but with a constraint on the shape of the time-dependence) (Picardi et al. 2021) or using continuous-time movement models (but with a complex implementation) (Hooten et al. 2014). Generally, we feel that ecologists are still lacking an easy and flexible tool to answer the question ‘when is an animal’s habitat selection changing?’, learning from the data with no assumption about when changes should happen.

Here, we present how dynamic logistic regression models allow one to easily estimate the temporal dynamics of habitat selection without *a priori* discretization of time. Dynamic logistic regression models are commonly used to analyze binary time series in survival analysis (Martinussen and Scheike 2006), but can be applied to other data sources (Fahrmeir 1992). First, we use simulated movement data to demonstrate that dynamic logistic regression models can adequately recover time-varying habitat selection coefficients. We also highlight the influence of parameters, whose values are under the researcher’s control, on the estimation process. Second, we illustrate the usefulness of the approach by applying time-varying HSA on blue wildebeest (*Connochaetes taurinus*) tracking data, showing how one can describe temporal variations of animal habitat selection such as diel, seasonal and post-release changes. The relevant R scripts are provided to facilitate the adoption of the method by ecologists.

## 2. Methods

### 2.1. Dynamic logistic models for time-varying HSA

#### 2.1.1. General principles of dynamic logistic models

Here, we briefly describe the discrete-time state space model developed by Fahrmeir (1992) to estimate time-varying coefficients from generalized linear models, and especially logistic models. Generally, discrete-time state space models relate observations over time to hidden parameters, with hidden parameters following a Markovian transition model (Auger-Méthé et al. 2021). Once applied to dynamic logistic regressions in the context of time-varying HSA, such relationships can be simplified as (1) and (2):

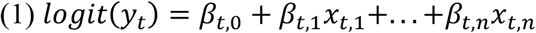

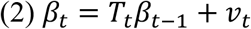

with *y*_*t*_ a binary response variable (i.e. used/available), *(β*_*t,1*_*, . . . , β*_*t,n*_) the hidden parameters (i.e. time-varying selection coefficient), *(x*_*t,1*_*, . . . , x*_*t,n*_) the covariates (i.e. environmental layers), *v*_*t*_ the error process with *v*_*t*_∼*N(*0, Q_*t*_), Q_*t*_ and *T*_*t*_ the covariance and transition matrices of the Markov chain at time *t*. The transition matrix *T*_*t*_ is estimated by the model. The covariance matrix Q_*t*_ is a diagonal matrix with initial values (at *t*_0_) chosen by the researcher, and then automatically updated over time during the fitting process. Hereafter, for simplicity, we call *Q* the initial value used by the researcher to fill in the Q_*t*_ diagonal matrix, and as it can greatly affect the wiggliness of the model (see Results), we also refer to *Q* as the wiggliness parameter. Time-varying coefficients *(β*_*t,1*_*, . . . , β*_*t,n*_) are estimated by maximization of the posterior densities, following a partial Bayesian approach. Basically, the generalized extended Kalman filter and smoother algorithm described by Fahrmeir and Kaufmann (1991) recursively iterates prediction, correction and smoothing steps to approximate posterior mode estimation in dynamic generalized linear models.

#### 2.1.2. Implementation in the context of HSA

To conduct time-varying HSA using dynamic logistic models, the following steps are required: first, as in RSF analyses, a sample of locations that could be considered ‘available’ is drawn. This can be, for instance, locations sampled randomly within the animal home range. Each used location (*y*_*t*_ = 1) obtained at time t is paired with *N* random locations (*y*_*t*_ = 0). Each used and available location is then characterized using environmental variables or any other variable of interest *(x*_*t,1*_*, . . . , x*_*t,n*_). Finally, the time-varying parameters *(β*_*t,1*_*, . . . , β*_*t,n*_) are estimated by fitting a dynamic logistic model, with type of location (used vs. available) as response variable *y*_*t*_ and the covariate time series *x*_*t,n*_ as predictors. Here, to fit the dynamic logistic model, we used the “ddhazard” function from the *dynamichazard* R package by (Christoffersen 2021), which implements the method described by (Fahrmeir 1992). We provide the R script needed to run a time-varying HSA on a simulated trajectory (Appendix S1).

### 2.2. Evaluation of the accuracy of dynamic logistic models for time-varying HSA

To assess the ability of our approach to detect shift in habitat selection patterns, we (1) simulated animal trajectories emerging from time-varying selection for one environmental variable, and (2) fitted dynamic logistic models on the simulated data to compare the estimated coefficients with the theoretical values used in the simulations.

#### 2.2.1. Landscape and movement simulation

For simplicity, animal trajectories were simulated on one habitat layer (500 × 500 cells), with values that did not vary over time. To mimic patchy landscapes, we used spatially correlated Gaussian random fields, which attribute a continuous value ranging from 0 to 1 to each cell, using the *localGibbs* R package (Michelot, Blackwell, and Matthiopoulos 2019). Following Michelot, Blackwell, and Matthiopoulos (2019), we then simulated animal trajectories over 500 time steps using a local Gibbs movement model. For each time *t*, 1000 potential locations were uniformly generated within a 100-pixel radius around the current location, and the location at time *t+1* was sampled among them with probabilities proportional to the strength of selection for potential locations. This strength of selection was determined by the value of the habitat layer at these locations, and by the model coefficient describing how the strength of selection changes with values of the habitat variable. An important benefit of using a local Gibbs model is that the coefficient of habitat selection used in the simulation model is theoretically equal to the one that should be estimated by a RSF fitted on the data (Michelot, Blackwell, and Matthiopoulos 2019).

#### 2.2.2. Scenarios of temporally-changing habitat selection

To be able to test whether changes in habitat selection strength could be robustly recovered by dynamic logistic models, we built scenarios that differed in terms of how often the model’s coefficient of habitat selection changed over time. We did so by randomly sampling, in the [-5, 5] range, and either every 20 steps (referred as “frequent change” scenario) or every 250 steps (referred as “rare change” scenario), the model’s habitat selection coefficient. To avoid having sudden, step-like, changes in habitat selection, we then used spline regressions to smooth the variations of habitat selection over time. For each scenario, we generated 100 trajectories per simulated landscape, and replicated this on 100 different landscapes. We then tested our ability to recover the temporal changes in the model’s habitat selection coefficient by fitting a dynamic logistic regression model to each trajectory as presented above, drawing 100 available locations, at each time-step, within the 99% utilization distribution location-based kernel of each simulated trajectory. We also assessed to what extent the model’s estimation was affected by the value of the wiggliness parameter *Q*. We did so by fitting, for each dataset of each scenario, a set of models with different values of *Q*, ranging from 0.01 (i.e. low wiggliness) to 2 (i.e. high wiggliness). For each value of *Q*, we then averaged the estimated coefficients over each set of 100 simulated trajectories per landscape, and fitted a linear regression with the mean estimated coefficient as response and theoretical coefficients which were used in the simulations as predictors, adding a random intercept with replication number. A slope near 1 would indicate that a dynamic logistic model is able to estimate the temporal changes in the habitat selection coefficient correctly, and an intercept near 0 would indicate that the estimations are not biased.

### 2.3. Time-varying HSA: applications

To illustrate some applications of time-varying HSA, we analyzed wildebeest movement datasets collected in Hluhluwe-iMfolozi Park (South Africa). There, surface-water availability, which is high during the wet season (October-March) as temporary waterholes are filled up by the rains, becomes low in the dry season (April to September), with water remaining available only in a few rivers. Since wildebeests are water-dependent grazers that preferentially forage in open grasslands, we used the distance to the closest main river and the habitat openness as relevant habitat variables to demonstrate the use of dynamic logistic models. For each study below, we fitted the models with a high-value for *Q* (Q=2), as simulations showed that high wiggliness in the estimates lead to better results (see Results section, or Appendix S2).

#### 2.3.1. Short-term temporal variations: diel changes in habitat selection

We used tracking data collected on one wildebeest over one month in the dry season, at a fix rate of one location every 15 minutes. Following the practical implementation of time-varying HSA described above, we estimated the habitat selection of this wildebeest by (1) generating, for each time t, 100 random locations within its home range, (2) extracting the environmental characteristics of used and random locations, and (3) fitting a dynamic logistic regression to compare the habitat openness and the distance to the closest main river between the used and available locations over time. A preliminary visual inspection of the spatial data suggested that day/night and longer, few-days, changes in locations occurred (Appendix S3: Figure S1). To check whether these changes were associated with changes in habitat selection, we estimated the temporal auto-correlation of the time-varying habitat selection coefficients.

#### 2.3.2. Long-term temporal variations: seasonal changes in habitat selection

We used tracking data collected on one wildebeest over one year, at a fix rate of one location every 15 minutes, and subsampled to one location per hour. A preliminary visual inspection of the spatial data showed that the individual moved mostly westward and then eastward during the period (Appendix S3: Figure S2). Hence, in addition to habitat openness and distance to the closest river, we added longitude to the model’s predictors. We generated 10 random locations, per time t, within its home range to fit the time-varying HSA. To provide an example of how one can subsequently delineate temporal periods that are homogeneous in terms of habitat selection, we further segmented the time series of each model’s coefficients, using the segmentation method described by (Patin et al. 2020) and implemented in the *segclust2d* R package.

#### 2.3.3. Event-based variations: post-release changes in habitat selection

We used the tracking data of three wildebeest simultaneously introduced in the park in October 2020. Data were collected over the 100 days following the release date, at a fix rate of one location every hour. Similarly to previous applications, we estimated wildebeest’s habitat selection drawing 100 random locations, for each time t, within their individual home ranges. To incorporate the dispersal of the released individuals into HSA, we added longitude and latitude to model’s predictors. Hence, we fitted a dynamic logistic model for each individual, using habitat openness, distance to the closest river, longitude and latitude as predictors.

## 3. Results

### 3.1. Theoretical evaluation

In general, we found that dynamic logistic models allowed us to adequately recover the temporal changes in the coefficients of habitat selection used in the simulations. When the true coefficients did not change often (‘rare change’ scenarios), the wiggliness parameter had little impact and estimations were always good (Fig. 1a-b; Appendix S2: Figure S1). When the true coefficients did change often (‘frequent change’ scenarios), it became critical to use high values of the wiggliness parameter to obtain estimates matching the theoretical coefficients (Fig. 1c-d; Appendix S2: Figure S1). Importantly, the effect of the wiggliness parameter tend to stabilize at large values of *Q* (Appendix S2: Figure S2), making it safe to use large values when investigating large and frequent changes in habitat selection.

**Figure 1.**
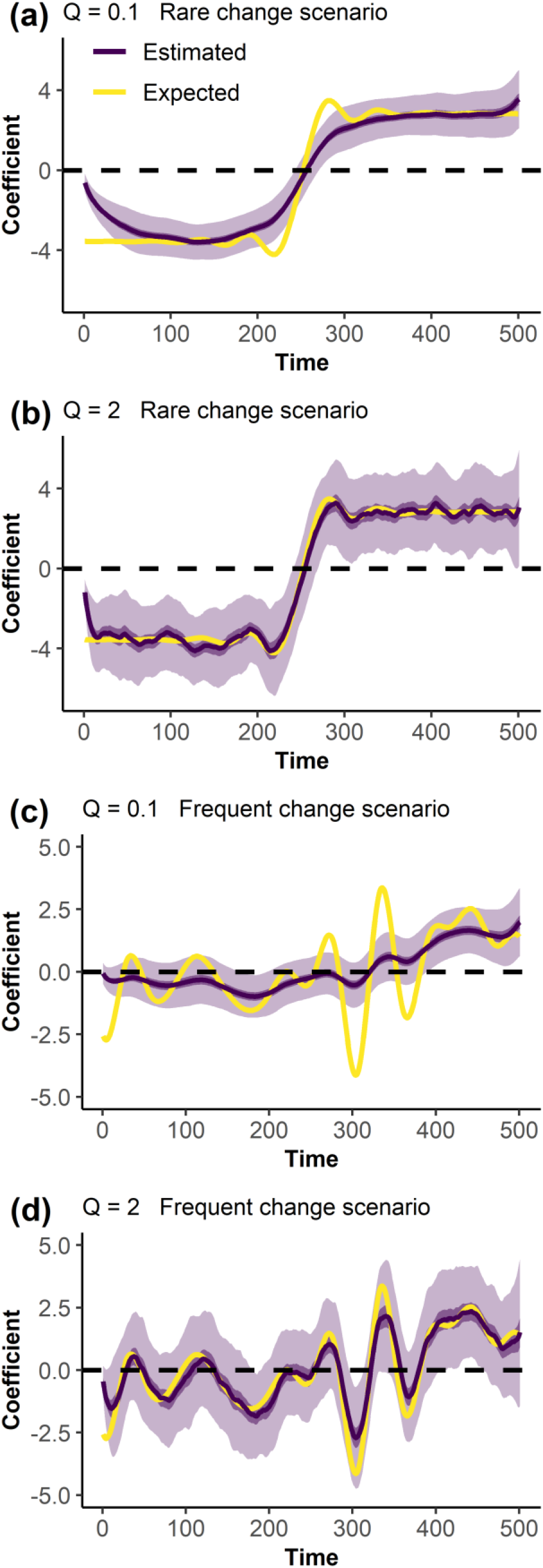
Estimated coefficient of habitat selection (purple) according to the frequency of change of the expected coefficient (yellow) and to the value of the model wiggliness parameter *Q*. The estimated coefficient is averaged on 100 simulations. Lighter ribbons show standard deviation, and darker ribbons show 95% confidence intervals.

### 3.2. Short-term temporal variations: diel changes in habitat selection

A time-varying HSA conducted with a dynamic logistic model very clearly revealed that the wildebeest’s selection for open habitats and rivers varied greatly over the month of the study (Fig. 2a-b). In particular, the wildebeest’s selection for open habitats clearly changed across the day/night cycle, as the auto-correlation period of the coefficient was approximately 24h (Appendix S3: Figure S3). Open habitats were strongly selected during nighttime, but were not selected, and sometimes even avoided, during daytime (Fig. 2a). We did not find such a pattern for the selection of areas close to rivers (Fig. 2b), but there were variations over periods of 3 or 4 days. Contrary to the diel variations in the selection of open habitats, such variations would be hard to detect using the common HSA approach based on time discretization.

**Figure 2.**
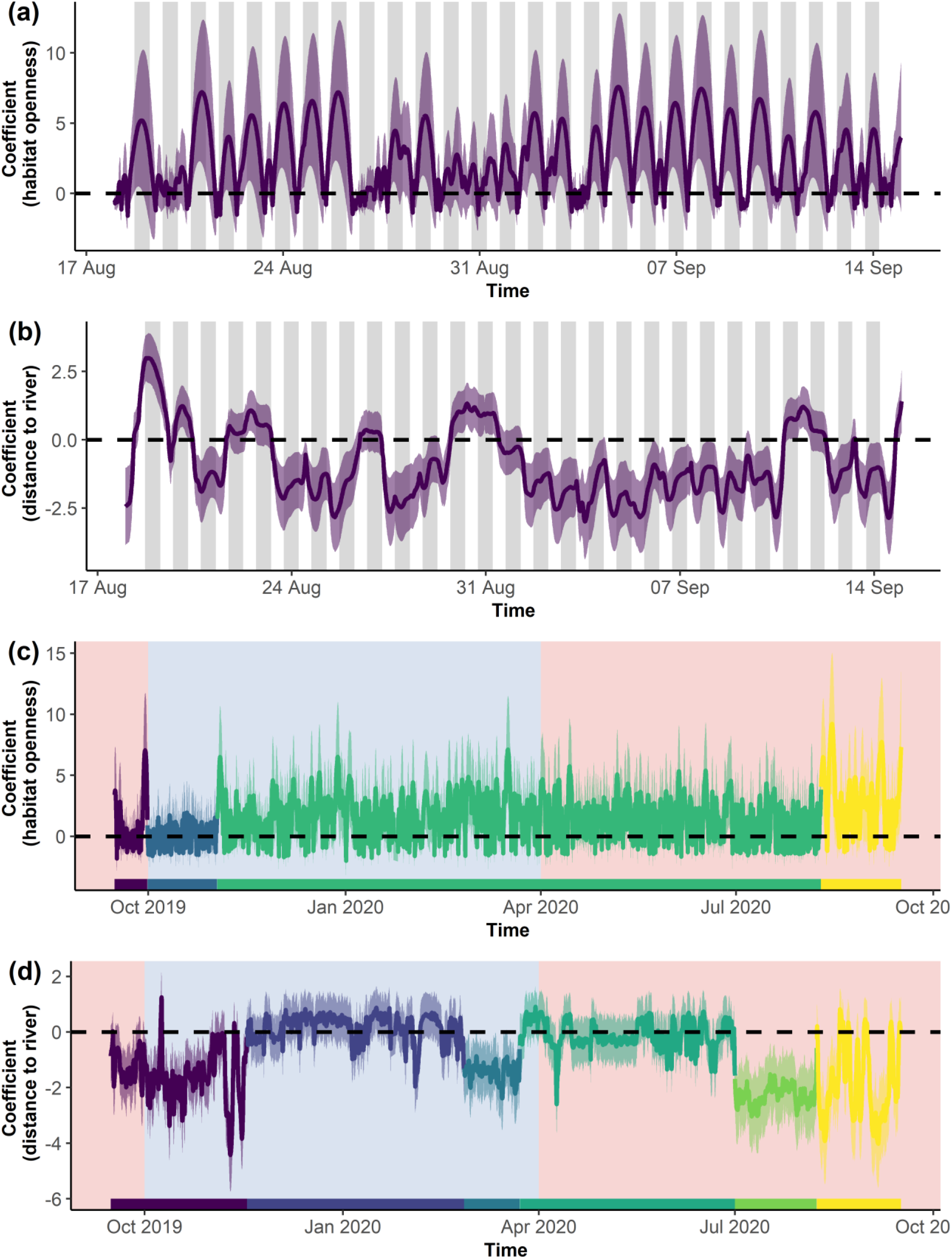
Estimations of short (a-b) and long-term (c-d) temporal variations of wildebeest’s selection for open habitats and areas close to rivers. Positive coefficients indicate selection for open habitats or areas far from rivers. Ribbons show 95% confidence interval. The night phases are shown by dark colors in panel (a-b). Common definitions of wet (blue) and dry (red) seasons are shown in background in panel (c-d), whereas segments of homogeneous habitat selection are colored on a purple-to-yellow gradient.

### 3.3. Long-term temporal variations: seasonal changes in habitat selection

The time-varying HSA showed clear seasonal changes in the wildebeest’ habitat selection, which could then be separated in several periods according to the segmented time-series (Fig. 2c-d). The existence and timing of some these periods were unpredictable *a priori*. For example, this wildebeest maintained the same overall strength of selection for open habitats from November to August, whereas this period covers months from both the dry and wet seasons (Fig. 2c). Also, we note that during the dry season (April to June) this wildebeest did not preferentially use areas close to rivers, but selected areas close to rivers consistently from July to mid-August (green segment) and likely did back-and-forth trips to and away from the rivers from mid-August to October (yellow segment) (Fig. 2d).

### 3.4. Event-based variations: post-release changes in habitat selection

After their release, the three wildebeest generally selected open habitats, but their level of selection differed between individuals (Appendix S3: Figure S4), particularly towards the end of the study period (Fig. 3a). Differences in the selection for areas close to rivers (Fig. 3b) or in the longitude (Fig. 3c) and latitude (Fig. 3d) of the park also became apparent at the end of the first month after release. Then, although the three wildebeest established in different areas (see difference in selection for longitude and latitude), the selection of areas close to rivers remained and was similar for two wildebeests (colored in green and yellow), whereas the third one avoided the areas close to rivers (colored in purple).

**Figure 3.**
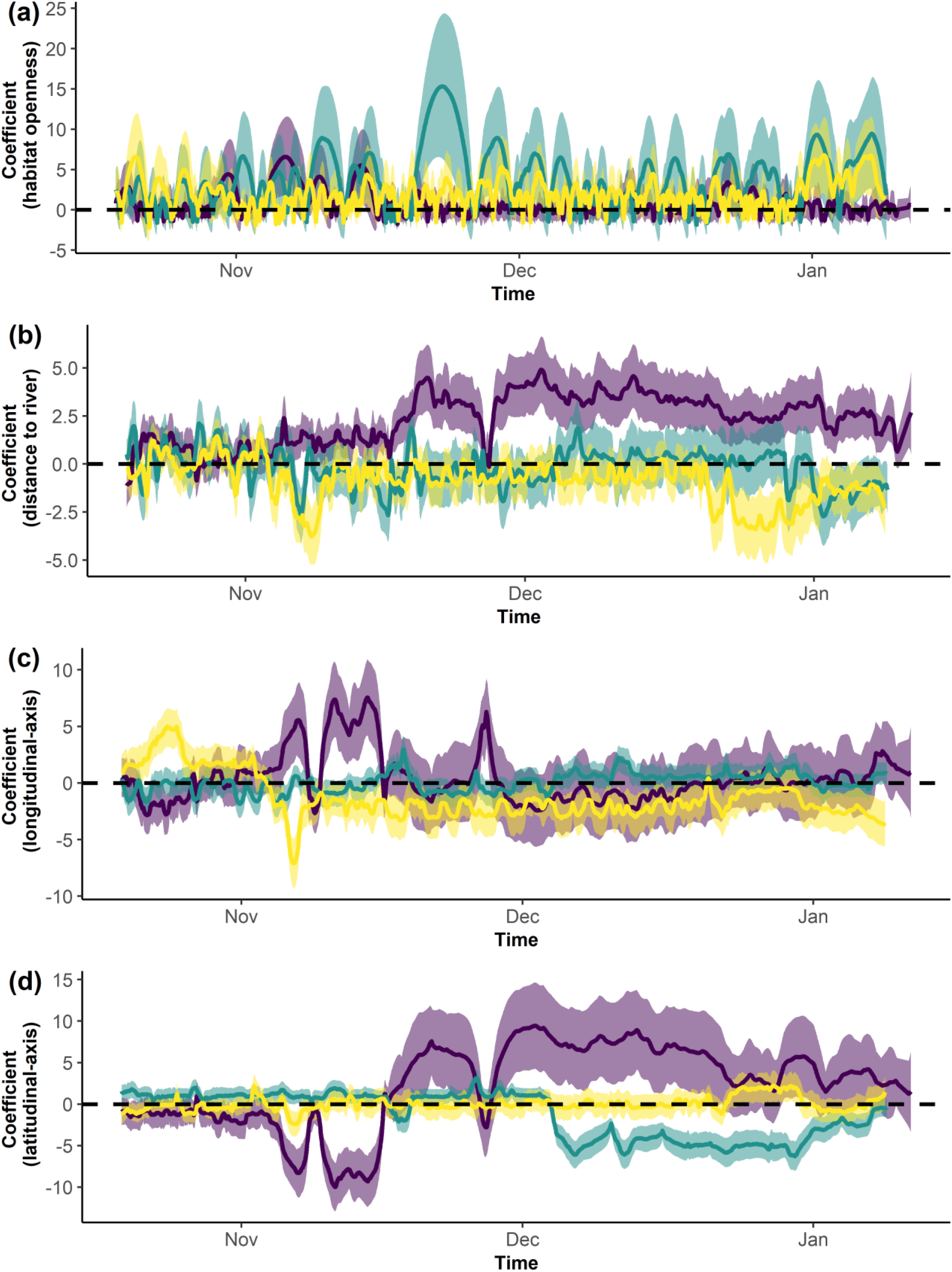
Estimations of post-release variation of wildebeest’s selection for open habitats, areas close to rivers, longitude and latitude. Positive coefficients indicate selection for open habitats, areas far from rivers, or areas at greater longitude and latitude. Estimated coefficients (line) and 95% confidence intervals (ribbons) are colored per individuals.

## 4. Discussion

There is obvious evidence that animals’ habitat selection changes regularly and at different time-scales, from diel to seasonal shifts, or during key life-history events such as dispersal Unfortunately, ecologists have had limited and often unsatisfactory options to study these changes. Most commonly, an *a prior*i discretization of time periods of apparent biological relevance is made, although this discretization can sometimes be difficult to justify, let alone to validate. The alternative approach of simply integrating time as predictor in a RSF has limitations (see discussion in Picardi et al. (2021)), and more statistically complex approaches (Hooten et al. 2014) are unlikely to be broadly used. In this work, we propose a new approach based on dynamic logistic models (Fahrmeir 1992) to easily estimate temporal changes in habitat selection, in a framework consistent with RSF. We demonstrated, using simulations, its general validity, while highlighting a point of attention (parameter *Q*). We also showcased its use for the study of diel, seasonal and post-release changes in habitat selection. In addition, we provided the necessary R scripts to make the use of time-varying HSA accessible to the ecologist community.

With this time-varying HSA approach, one can simultaneously estimate both the timing and the amplitude of habitat selection changes. Estimation of the timing of change from the data is what makes this approach novel and attractive. Many times, *a priori* discretization of time is a guesstimate from expert knowledge or is based on ancillary data (e.g. climate data) whose relevance for a particular dataset is not warranted. ‘Letting the data speak’ allows revealing the actual pattern of change. This may be of particular importance, for instance, in the study of inter-individual variability, as the timing of change can be one of the differencing variables, as evidenced in our post-release study case. As recognized by Picardi et al. (2021), time-varying HSA opens a new avenue to broaden the scope of the studies of inter-individual differences in space use, which has so far focused on movement characteristics or habitat selection strength. More generally, even when the relevance of an *a priori* discretization of time is easier to ascertain, such as when comparing daytime to night-time habitat selection, time-varying HSA allows one to immediately identify unusual periods (e.g. night of the 27^th^ of August, when the wildebeest did not increase its selection for open habitat). These unusual periods may either be of interest (in such case, one would have to conduct one standard HSA per night to have discovered this), or be ‘noise’ that should not affect the estimation of habitat selection strength during more ‘usual’ periods (conversely to what occurs in a standard HSA).

Importantly, data-driven estimation of the timing of change in habitat selection makes it possible to derive ‘segments’ of homogeneous habitat selection, and open the way to estimate specific habitat selection ‘modes’ of animals defined by the strength of habitat selection. Until now, behavioral modes in space use could be defined by movement characteristics such as speed and turning angles (Patin et al. 2020; McClintock and Michelot 2018), but could not integrate information about habitat selection, although it was likely to strongly contribute to define these modes. The use of segmentation-clustering algorithm such as *segclust2d* or hidden-Markov models on the time-series coefficients obtained by time-varying HSA will allow one to do this, and to estimate the duration and frequencies of such modes. This could complement very recent works developing behavioral-mode detection approaches based on step-selection functions (Prima et al. 2022; Klappstein, Thomas, and Michelot 2022).

Results from the time-varying approach proposed here are to some extent sensitive to the model’s wiggliness. In particular, but without surprise, model allowing for little wiggliness (small values of *Q*) can provide a poor fit to the data when habitat selection often changes. Models allowing for high wiggliness generally performed much better, especially if habitat selection often changed. There was no obvious evidence of an optimum value of *Q* to look for, as correlations plateaued when increasing *Q* values. Therefore, running a time-varying HSA with a high value for *Q* appears a safe way to conduct robust analyses.

In conclusion, we think dynamic logistic models offer an easy yet powerful approach to conduct time-varying HSA, for both exploratory and inferential studies. Our work, in which we show real-world applications and provide R scripts, aims to facilitate the appropriation of the method by ecologists and enriches their statistical toolbox. Novel questions about how animals time their response to environmental changes can now be addressed.

## Acknowledgements

We are grateful to Ezemvelo for providing the authorization to conduct this research, and to its ecologist D.J. Druce for his continuous administrative and field support, as well as for a review of the manuscript. We acknowledge the field efforts of B. Gehr and R. Leeming and anyone involved in the wildebeest captures. G. Fradin created the layer for habitat openness. This work was partially funded by the French National Research Agency (program ANR-16-IDEX-0006) through the i-Site MUSE (project REPOS).

## Supplementary information

## Appendix S1

Available in the figshare digital repository: https://doi.org/10.6084/m9.figshare.c.6365415.v3

## Appendix S2 Computer simulations to evaluate time-varying HSA

**Figure S1.**
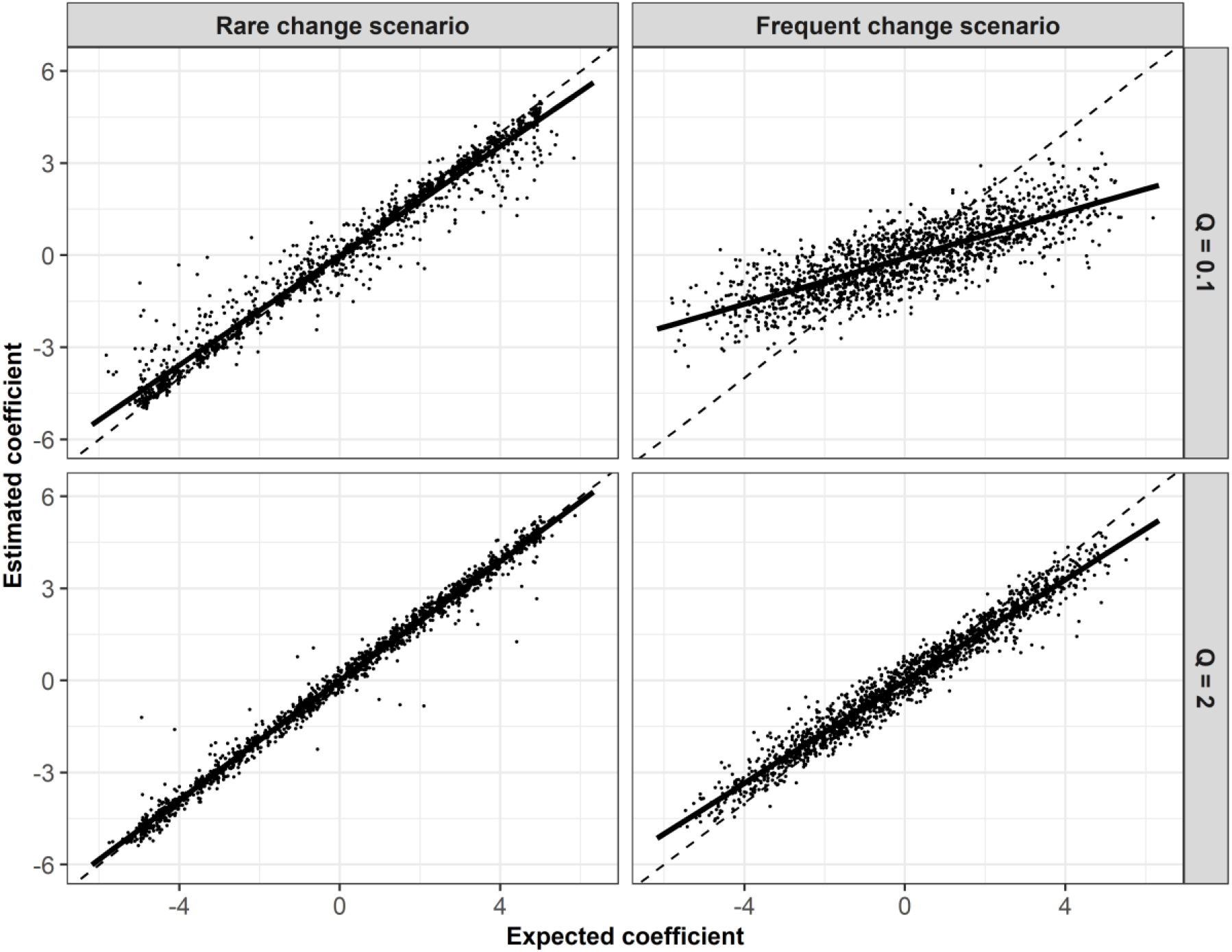
Comparison between the estimated and expected coefficients of habitat selection, according to the value of the wiggliness parameter Q used in the dynamic logistic model. High-wiggliness models performed better than low-wiggliness models to accurately estimate time-varying coefficients of habitat selection, both when habitat selection changes rarely and frequently over time.

**Figure S2.**
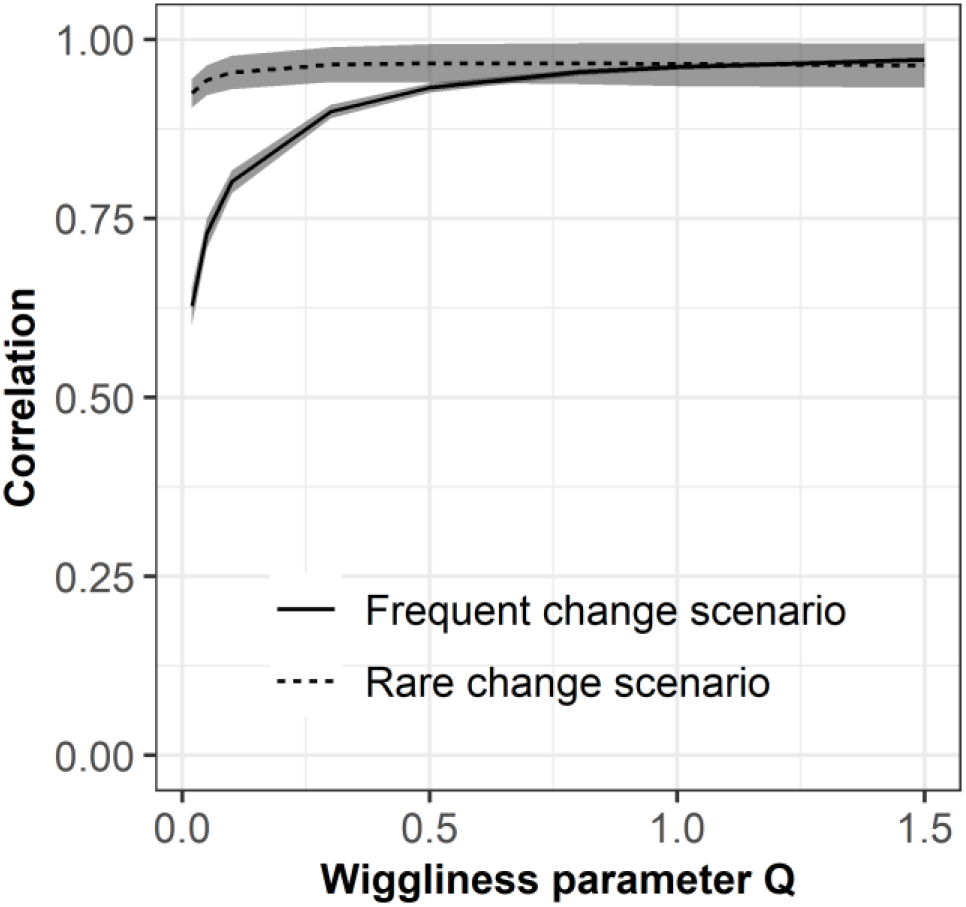
Relationship between the value of the model wiggliness parameter Q and the correlation between (1) the expected time-varying selection strength and (2) the average time-varying selection strength estimated on 100 replicates on 40 different landscapes, according to the frequency of change in habitat selection. Time-varying HSA was not sensitive to the choice of model wiggliness to estimate rare changes of habitat selection. On the contrary, high-wiggliness models performed better than low-wiggliness models to estimate frequent changes of habitat selection.

## Appendix S3 Applications of time-varying HSA on wildebeest examples

**Figure S1.**
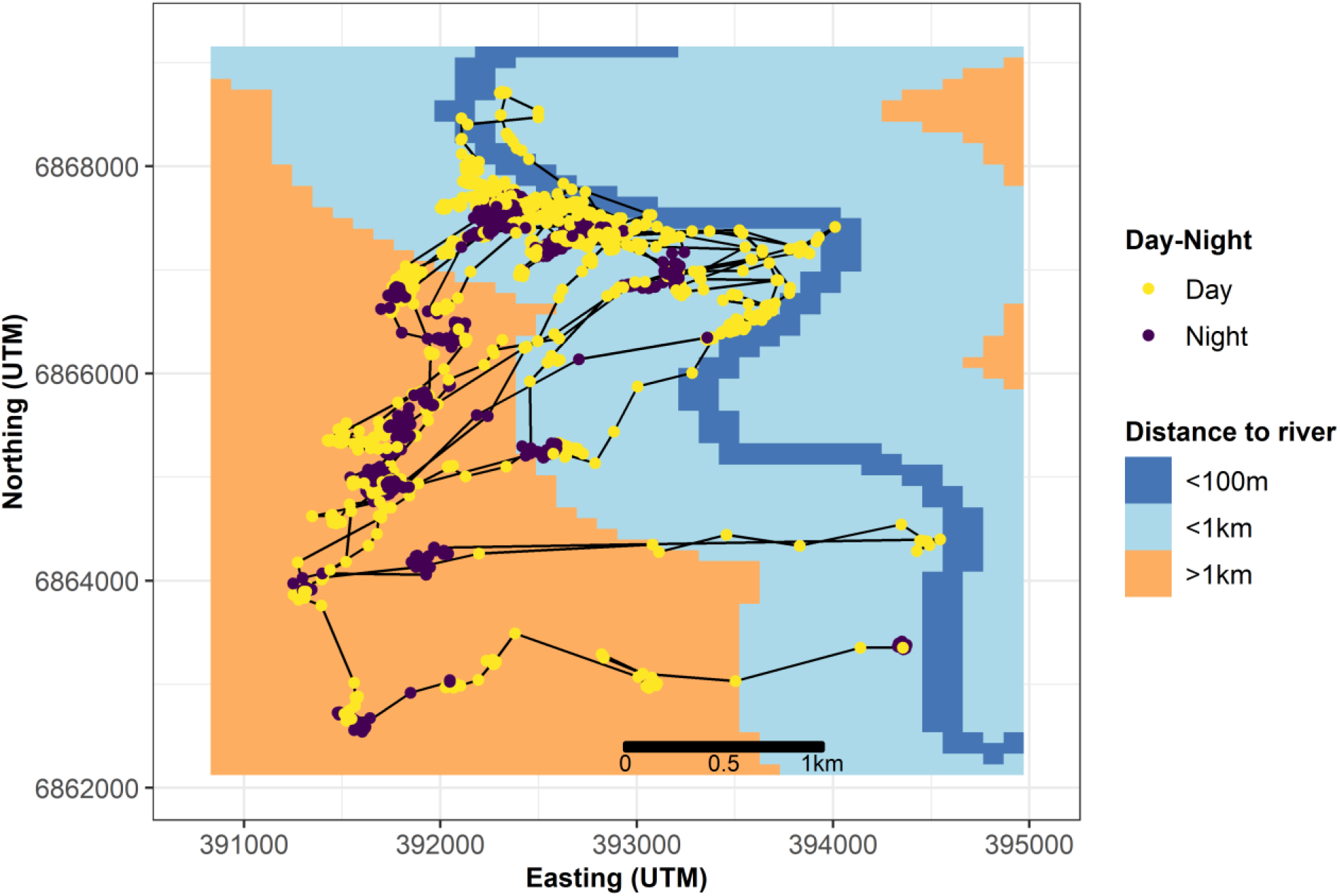
Illustration of the wildebeest’s locations recorded during one month in the dry season, at a fix rate of one location every 15 minutes. Background shows the distance to the closest river. Locations collected during the night are colored in purple, whereas those collected during the day are in yellow.

**Figure S2.**
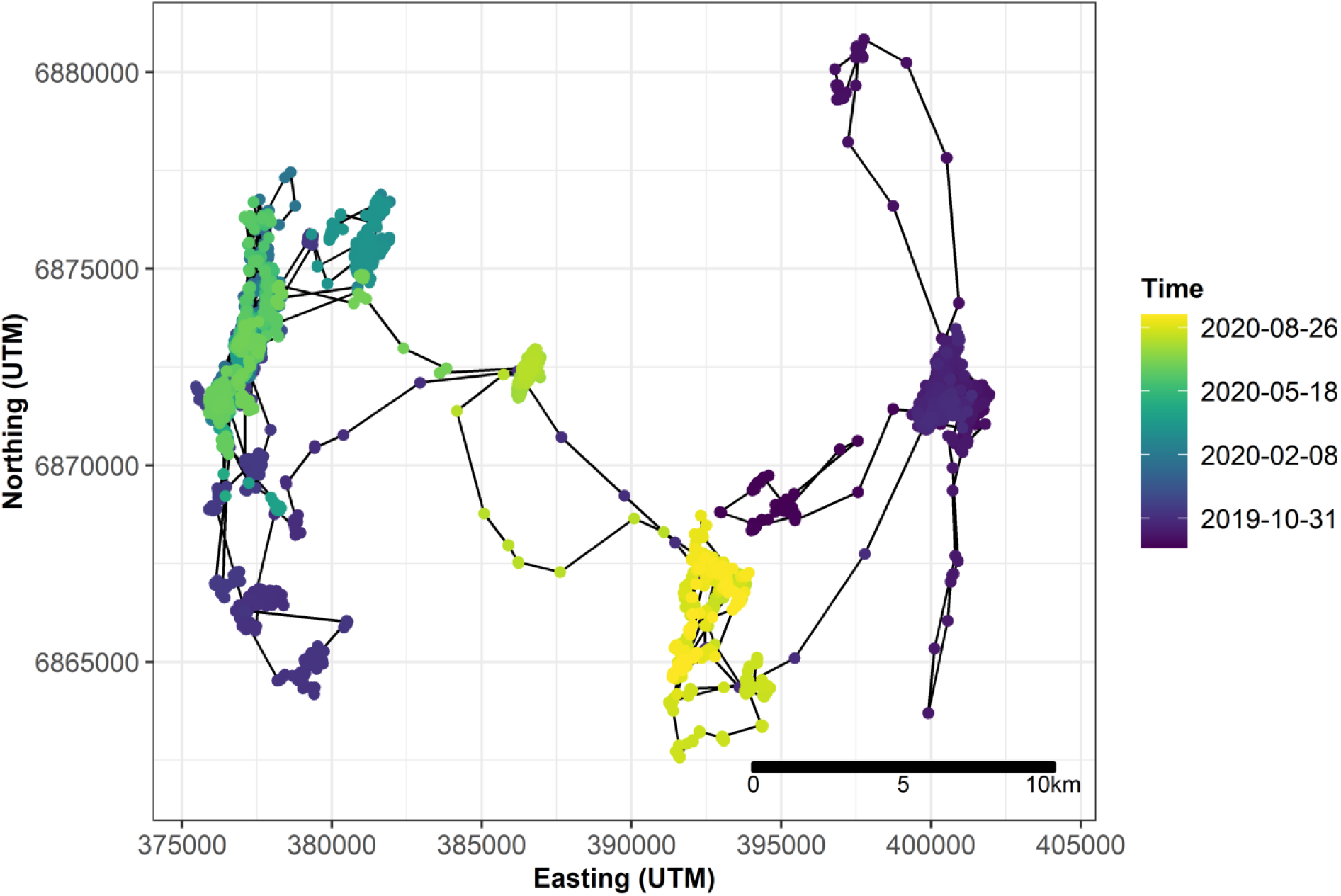
Illustration of the wildebeest’s locations recorded during one year, at a fix rate of one location every 15 minutes and subsampled to one location per hour. Locations are colored based on their collection time.

**Figure S3.**
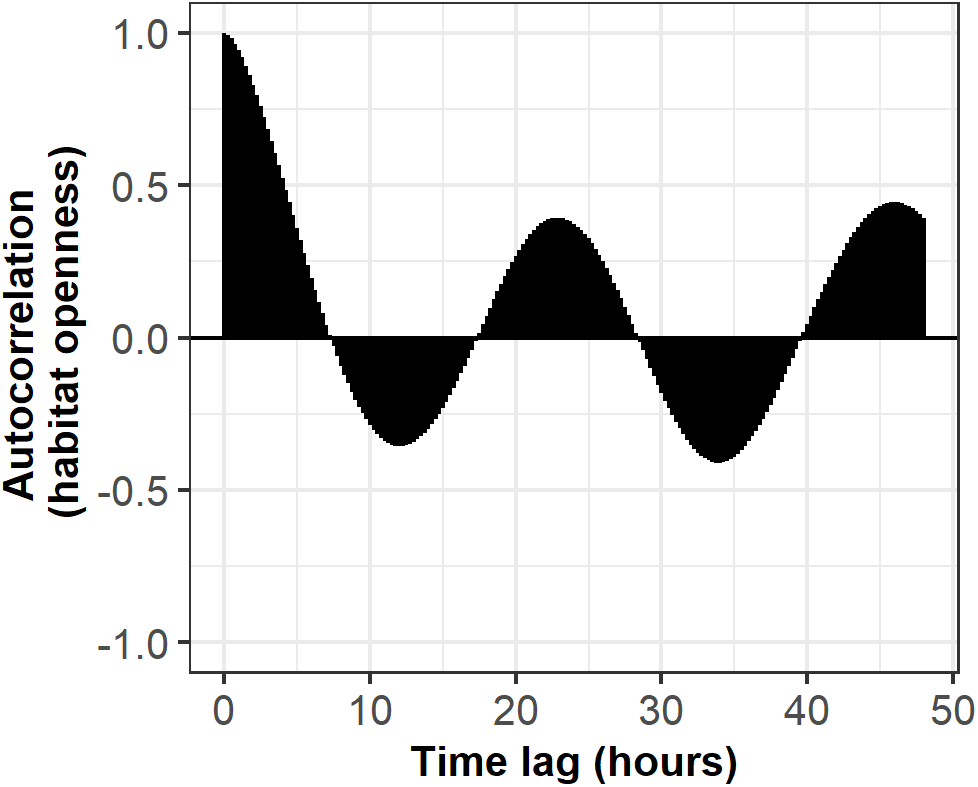
Temporal autocorrelation of the selection coefficient for open habitat *β(t)β(t* + *lag)*. Values close to 1 (resp. −1) indicate high auto-correlation, with similar selection strength (resp. opposite selection strengths), while values close to 0 indicate low auto-correlation.

**Figure S4.**
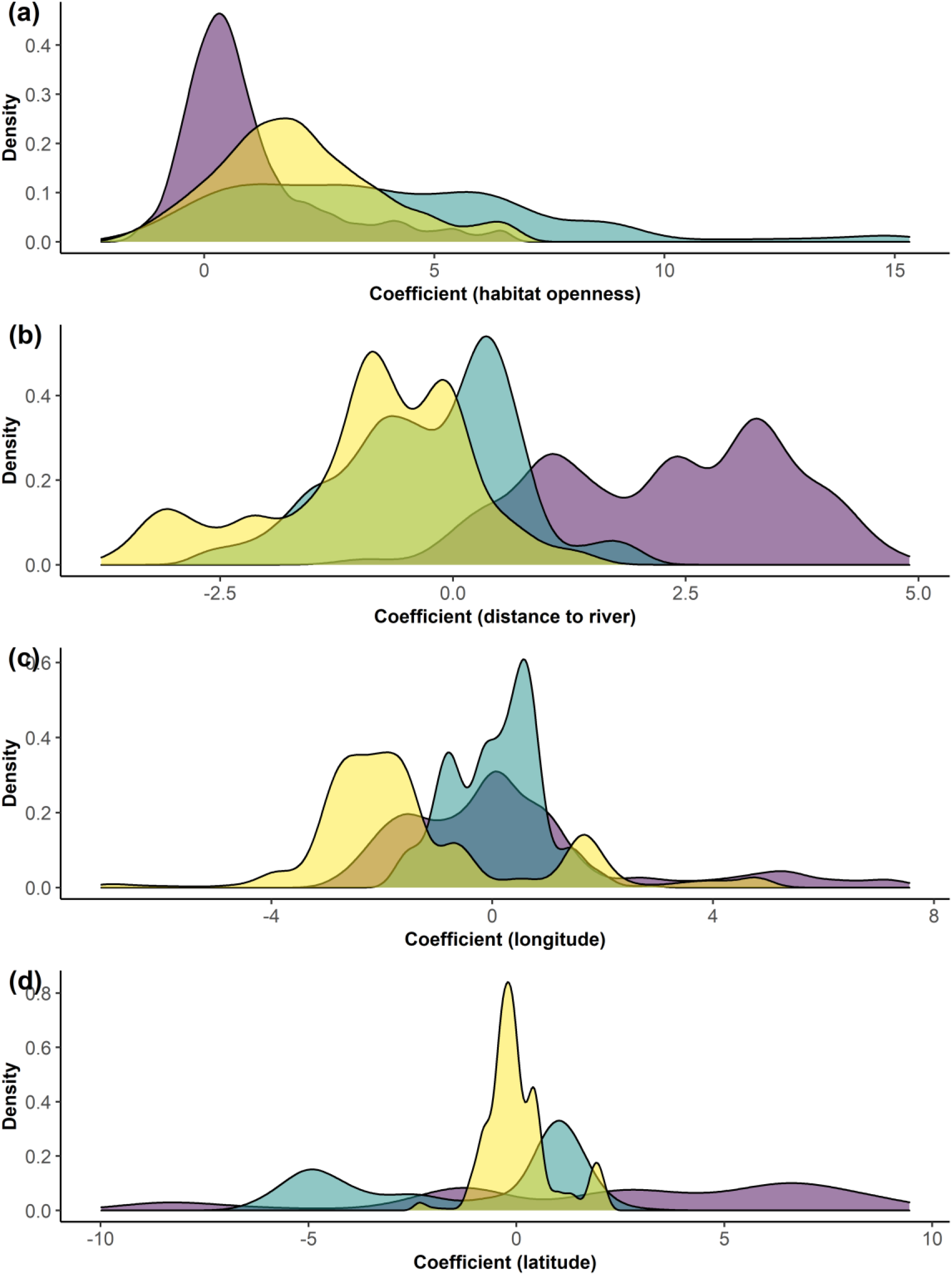
Distribution of the habitat selection coefficients for (a) habitat openness, (b) distance to the closest river, (c) longitude and (d) latitude. Each color shows the distribution of the selection coefficients for one individual.

